# ESI mutagenesis: A one-step method for introducing point mutations into bacterial artificial chromosome transgenes

**DOI:** 10.1101/844282

**Authors:** Arnaud Rondelet, Andrei Pozniakovsky, Marit Leuschner, Ina Poser, Andrea Ssykor, Julian Berlitz, Nadine Schmidt, Anthony A Hyman, Alexander W Bird

## Abstract

Bacterial artificial chromosome (BAC)-based transgenes have emerged as a powerful tool for controlled and conditional interrogation of protein function in higher eukaryotes. While homologous recombination-based recombineering methods have streamlined the efficient integration of protein tags onto BAC transgenes, generating precise point mutations has remained less efficient and time-consuming. Here we present a simplified method for inserting point mutations into BAC transgenes requiring a single recombineering step followed by antibiotic selection. This technique, which we call ESI (**E**xogenous/**S**ynthetic **I**ntronization) mutagenesis, relies on co-integration of a mutation of interest along with a selectable marker gene, the latter of which is harboured in an artificial intron adjacent to the mutation site. Cell lines generated from ESI-mutated BACs express the transgenes equivalently to the endogenous gene, and all cells efficiently splice out the synthetic intron. Thus, ESI-mutagenesis provides a robust and effective single-step method with high precision and high efficiency for mutating BAC transgenes.

## Introduction

The ability to precisely query functional hypotheses of protein function in cells requires the capacity to express rationally mutated proteins from genes under their native physiological regulation. Traditionally, transgenes expressed in higher eukaryotes are derived from cDNAs, and thus lack native cis-regulatory elements or alternative splicing isoforms, often resulting in overexpression and deregulation artefacts. This may hinder proper phenotypic functional characterization of mutated genes, as well as the determination of the precise localization and interaction partners of the protein products. While the development of sequence-specific nucleases has enabled mutation of specific genes at their endogenous loci(Gaj *et al.*, 2013; Ran *et al.*, 2013), mutations in essential genes that are lethal will prevent growth and recovery of viable cells. Additionally, deleterious mutations are prone to accumulate suppressive changes in chromosome integrity or gene expression during the procedure of selecting and expanding cells for analysis, particularly in the genomically instable cancer cell lines frequently employed.

The use of bacterial artificial chromosomes (BACs) as transgenes largely overcomes these limitations. BACs are bacterial vectors containing fragments of an eukaryotic genome that are large enough to encode an entire gene in its genomic context, including *cis*-regulatory elements. Transgenes expressed from BACs thus deliver near-physiological expression in eukaryotic cells(Kittler *et al.*, 2005; Bird and Hyman, 2008; Poser *et al.*, 2008; Bird *et al.*, 2011). BAC transgenes modified to be RNAi-resistant allow for conditional exposure of recessive mutations through selective depletion of endogenous protein(Kittler *et al.*, 2005; Bird and Hyman, 2008; Ding *et al.*, 2009; Bird *et al.*, 2011; Scolz *et al.*, 2012; Zheng *et al.*, 2014; Rondelet *et al.*, 2020).

Because of their size, BACs do not lend themselves to restriction/ligation-based modification techniques. Instead, homologous recombination-based recombineering techniques are used to engineer BACs in *Escherichia coli* (Murphy, 1998; Zhang *et al.*, 1998; Yu *et al.*, 2000). Recombineering utilizes phage proteins (Redα, β and γ from phage λ; or RecE and RecT from the Rac prophage) to promote homologous recombination, facilitating a wide variety of DNA modifications. The high efficiency of the single-step recombineering procedure to insert a protein tag attached to a selectable marker gene on a DNA vector has allowed its adaptation to genome-scale high-throughput pipelines(Sarov *et al.*, 2006; Poser *et al.*, 2008; Sarov *et al.*, 2012; Hasse *et al.*, 2016; Sarov *et al.*, 2016). The subsequent generation of transgenic cell lines based on these libraries has further been used to precisely determine the cellular localisation and the quantitative interactome of more than a thousand proteins (Hubner *et al.*, 2010; Hutchins *et al.*, 2010; Hein *et al.*, 2015). On the contrary, current techniques to introduce point mutations in BACs still require either a counterselection-based two-step procedure(Bird *et al.*, 2011; Wang *et al.*, 2013) or a lower efficiency one-step procedure requiring extensive PCR-screening (Lyozin *et al.*, 2014). Here we present a simple and efficient one-step procedure to introduce point mutations in BAC transgenes, harnessing introns to carry selectable markers, which reduces the time and cost of generating mutagenized constructs.

Modifying intronic sequences is an attractive approach to modifying or gaining gene functionality with minimal perturbation of the host gene protein product, and has met use in a variety of applications such as conditional knock-out and mutant constructs, or miRNA expression(Gu *et al.*, 1994; Kaulich *et al.*, 2015; Wassef *et al.*, 2017). Synthetic introns have been designed based on the four core splicing signals necessary for spliceosome processing: two splice sites (SS) located at the 5’ and 3’ intron boundaries (5’SS and 3’SS, respectively), a branchpoint sequence (BP) located approximately 25 nucleotides upstream of the 3’SS, and a polypirimidine tract (PPT) directly upstream of the 3’SS (Figure 1A) (Lin *et al.*, 2003; Wang and Burge, 2008; Mercer *et al.*, 2015). Most of the sequence determinants for splicing are thus located within the intron. In most eukaryotes, the consensus sequences of the core splicing elements located within exons are limited to a (C/A) A G sequence directly upstream of the intron and a G downstream of the intron (Figure 1 A) (Zhang, 1998). Synthetic introns designed from these minimal core splicing signals are indeed efficiently spliced out of an exon, and have been used in multiple applications, such as making conditional knockouts, or expressing shRNAs or miRNAs(Lin *et al.*, 2003; Greber and Fussenegger, 2007; Lin and Ying, 2013; Seyhan, 2016; Guzzardo *et al.*, 2017).

**Figure 1:**
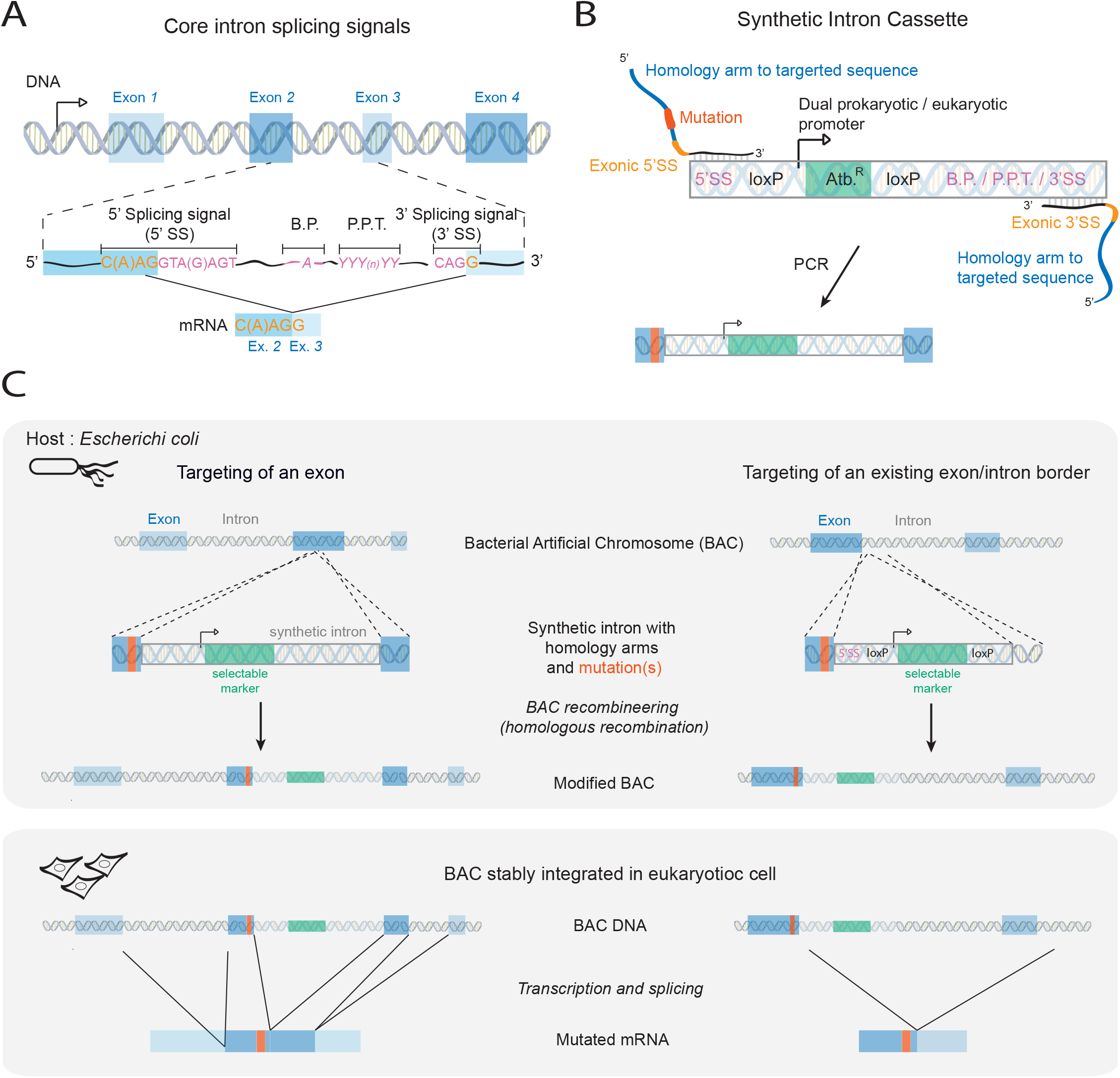
ESI-mutagenesis: a one step recombineering procedure to introduce point mutations in BAC. **(A)** Scheme showing the organisation of an eukaryotic gene with exons (in blue) and their flanking introns. Introns are spliced out from pre-messenger RNAs to produce mature messenger RNAs (mRNAs) containing only exons. The core splicing signals constitute the minimal information required for the splicing of an intron, and include a 5’ and 3’ splicing signal sequences (5’SS and 3’SS, respectively), a branch point (B.P.), and a poly-pyrimidine tract (P.P.T). The position of these signals relative to the exon / intron border is indicated with the corresponding nucleotides shown in pink (in introns) or light orange (in exons). ESI mutagenesis is a one step recombineering procedure that relies on the introduction into a BAC of a synthetic intron coding a selectable marker along with the mutation of interest, thereby allowing for the easy selection of correct recombinants. After transcription of the transgene in eukaryotic cells, the synthetic intron is spliced out to produce a mutated mRNA. **(B)** The synthetic intron constitutes of the intronic core splicing signal (in pink), an antibiotic resistance cassette (Atb.^R^) under the control of a dual eukaryotic/prokaryotic promoter (in green), and two loxP sites flanking the antibiotic resistance cassette. In order to serve as a template for BAC recombineering, the synthetic intron is amplified by PCR with primers containing ~50 bp long homology arms to the targeted sequence (in blue), the mutation(s) to introduce (in orange) and the exonic part of the 5’SS and 3’SS (in light orange). **(C)** Scheme showing the introduction of a point mutation into a BAC by ESI-mutagenesis. Depending on the position of the sequence to mutate, the point mutation can be inserted along with the synthetic intron into an exon (Left panel), or, if the mutation is located in the proximity of a pre-existing intron, the part of the synthetic intron encoding the antibiotic resistance can be targeted into this intron (Right panel). The synthetic intron is spliced out during RNA maturation and only the desired mutation is present in the mRNA.

Here we present a one-step BAC recombineering procedure in which point mutations are introduced into a BAC along with a synthetic intron. An antibiotic-resistance gene located within the intron allows the selection of positive clones after the single recombineering step in bacteria, and is seamlessly spliced out of mRNA sequences of the harbouring transgene in eukaryotic cells. This procedure significantly increases the efficiency and decreases the screening time of BAC mutagenesis.

## RESULTS

### ESI-mutagenesis: a one-step BAC recombineering strategy for introduction of point mutations

The ESI-mutagenesis strategy is outlined in Figure 1. First, a synthetic intron “cassette” is amplified by PCR for use in the recombineering reaction (Fig 1B). This synthetic intron cassette contains the core intron-based splicing signal sequences, as well as an antibiotic resistance gene. We have created multiple cassettes with varying bacterial and eukaryotic selectable markers, presented in Figure S1. In order to become a functional intron, the synthetic intron cassette must be recombined into a site in the exon where it is directly bordered by an upstream (C/A) A G sequence and a downstream G (Figure 1A, and C). While this (C/A) A G | G sequence would occur every 128 nucleotides assuming random distribution, similar sequences can be mutated to this sequence at amino acid wobble positions during cassette amplification (see Figure S2 for an example), significantly increasing the frequency of useable integration sites. Furthermore, this consensus is not a strict requirement, and a variety of sequences are found in cells and predicted to function(Zhang, 1998; Nguyen *et al.*, 2018; Ohno *et al.*, 2018). Therefore, it is unlikely that a potential synthetic intron integration site will not be found within the useable vicinity of a mutation site of interest. In addition, if the desired mutation is close enough to an existing intron, the antibiotic resistance cassette can be directly targeted to the endogenous intron by generating only one new splice site, with the opposite homology arms targeting intronic sequences (see Figure 1C, right side). This is preferable when possible, because not only does it maintain more closely the native intron/exon structure, but it also avoids the creation of unusually small exons (less than ~50bp), which may be prone to exon skipping (Dominski and Kole, 1991).

An example of how to design ESI-mutagenesis is presented in supplementary Figure S2. Once a target site for synthetic intron insertion near the mutation site is identified, primers are designed to amplify the synthetic intron cassette and containing 5’-extended 50 bp arms homologous to sequences directly surrounding the insertion site. One or both homology arms contain the mutation(s) to be integrated. The cassette is amplified by PCR and the resulting product is inserted into the target BAC via recombineering (Figure 1 C). Correct recombination events are selected using the antibiotic resistance encoded within the synthetic intron. A protocol for the ESI mutagenesis recombineering procedure as well as generating mammalian transgenic cell lines is provided in Figure S3.

Upon transfection of the modified BAC into eukaryotic cells of interest, cells that have integrated the BAC in their genome may also be selected using the antibiotic resistance encoded within the synthetic intron. If desired, the antibiotic resistance gene may be removed by application of Cre recombinase via loxP sites located within the cassette (Figure S1). Once the BAC transgene is transcribed the synthetic intron is spliced out leading to the formation of a messenger RNA carrying the desired mutation.

### ESI mutated BAC transgenes yield proteins with the expected size and localization

To confirm that BAC transgenes containing synthetic introns yielded proteins of the correct size and localisation when expressed in human cells, we started with a panel of 10 GFP-tagged BAC transgenes, and modified each using our ESI-mutagenesis strategy to introduce mutations to render them resistant to specific siRNA targeting sequences (Table S1). We then generated Hela cell lines stably expressing either the “parental” GFP-tagged genes, or the “ESI-mutated” (i.e. RNAi-resistant) GFP-tagged genes (Figure 2A). Because these mutations should not change the corresponding protein sequence, this allowed us to directly compare the expression and localization of the mutated transgenes to that of the parental, while offering an assay for the presence and function of the introduced mutation (RNAi resistance). Depending on the location of the mutation with respect to an existing intron, we either targeted the synthetic intron along with the desired mutation into an exon (4/10 BACs) (Figure 1 C left panel, Figure 2B), or made use of an existing exon-intron border to introduce the desired mutation and the antibiotic resistance (6/10 BACs) (Figure 1 C right panel, Figure 2B). All stably integrated “ESI-mutated” transgenes led to the expression of proteins of sizes identical to the one expressed from the corresponding parental transgenes (Figure 2 C). Moreover, we could not observe any differences in the cellular localisation of proteins expressed from “ESI-mutated” transgenes as compared to those expressed from parental transgenes (Figure 2 D). We next performed siRNA experiments on six of the parental / “ESI-mutated” BAC lines pairs to confirm functionality of the mutations. When the parental BAC-GFP lines were treated with the siRNA against the tagged transgene, we observed a reduction in the corresponding mRNA levels comparable to that observed in Hela wild type cells (Figure 3 A). In contrast, ESI-mutated BAC lines did not show any reduction in the corresponding mRNA levels upon siRNA treatment. We then analysed RNAi-resistance on the protein level for two of the ESI-mutated BAC lines for which we had the corresponding specific antibody. Both CEP135 and AURKB ESI-mutated BAC clonal lines showed expression of their respective GFP-tagged transgenes at a level similar to the corresponding endogenous proteins (Figure 3 B), and treatment of both ESI-mutated lines with the corresponding siRNA resulted in depletion of the endogenous protein while the transgene could still be detected (Figure 3 B).

**Figure 2:**
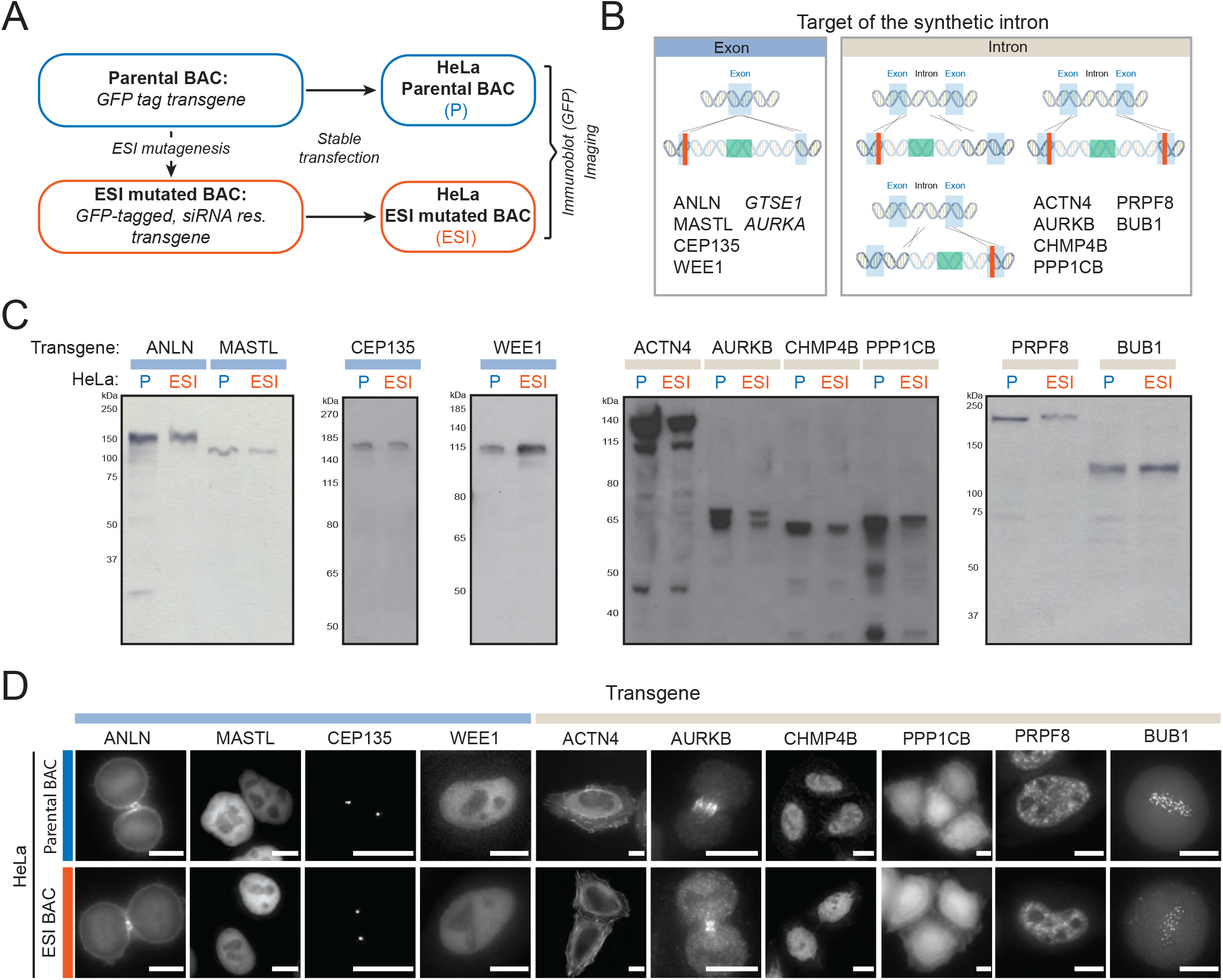
ESI-mutated BAC transgenes yield proteins of the right size and localisation. **(A)** GFP tagged BAC transgenes (Parental BAC, P) were mutated by ESI mutagenesis to be RNAi resistant (ESI mutated BAC, ESI) and transfected into HeLa cells. **(B)** Depending on the position of the sequence to mutate, the synthetic intron was either targeted into an exon (left panel, light blue label in B, C and D) or into the neighbouring intron by making use of pre-existing 5’SS, and / or 3’SS (right panel, light brown label in B, C and D). Names of the mutated transgene are indicated in the panel corresponding to their mutation strategy. The ESI mutated AURKA transgene is analysed in Figure 4. The GTSE1 transgene was ESI mutated at its SxIP motifs and is analysed in Figure 3. **(C)** GFP-tagged BAC transgenes ESI mutated to be RNAi resistant yield proteins of the expected size. Immunoblotting on cell-lysate of pools of HeLa cells transfected with either parental BACs (P) or ESI-mutated BACs (ESI). GFP antibody was used as a probe. Transgenes names are indicated at the top. **(D)** GFP-tagged transgenes ESI mutated to be RNAi resistant show the same cellular localization as their parental GFP-tagged transgenes. Still images of live cell imaging on HeLa stably transfected with the parental or the ESI-mutated BAC. CHMP4B and AurkB -GFP transgenes were detected by immunofluorescence with anti-GFP antibody. Because of differences in expression level within cell pool, each picture was acquired and scaled independently of the others. Scale Bar 10 μm.

**Figure 3:**
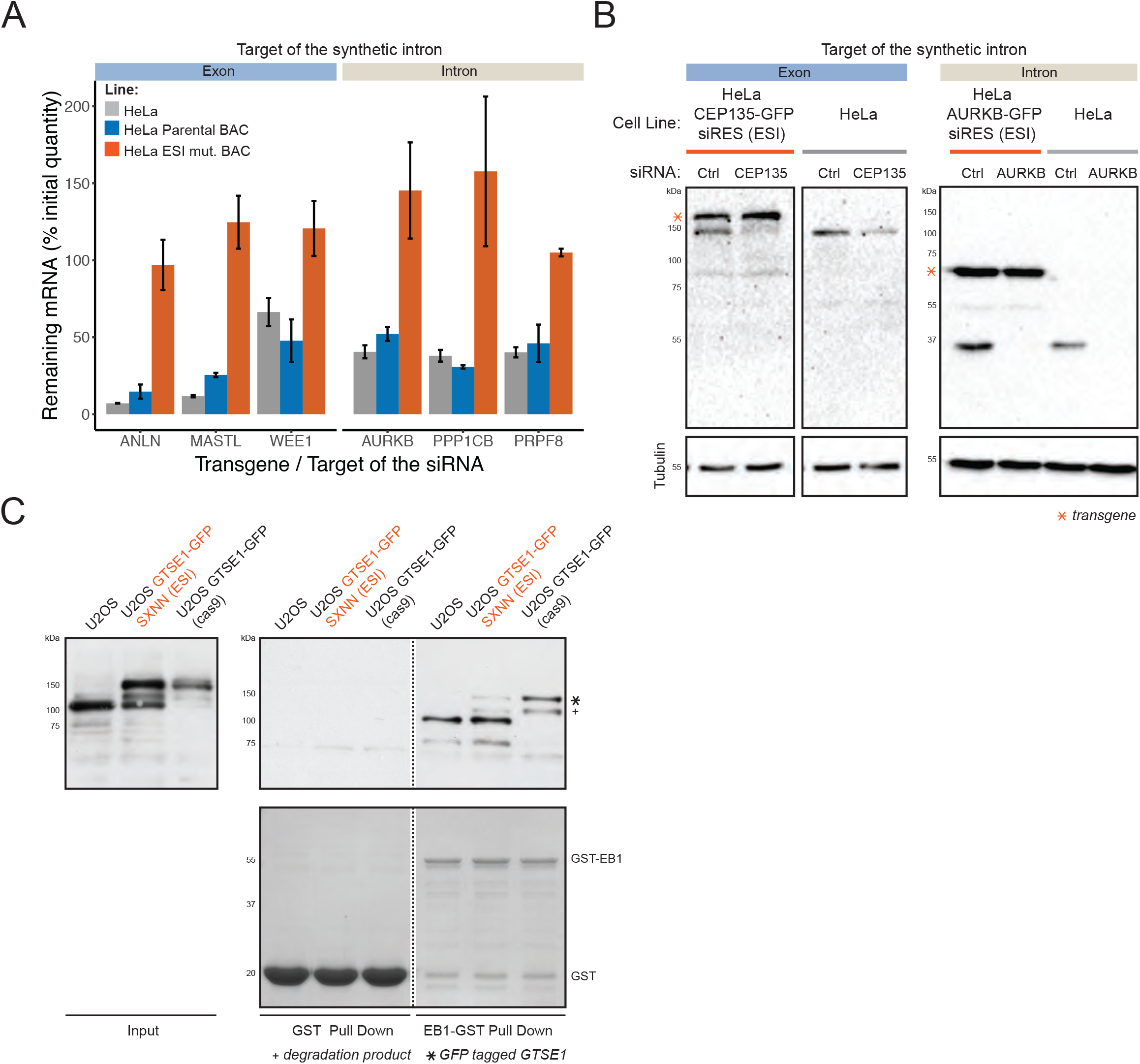
ESI-mutated BAC transgenes show the expected phenotype. **(A)** BAC transgenes ESI mutated to be RNAi resistant show RNAi resistance at the mRNA level. HeLa wild type, parental BAC lines, and ESI-mutated BAC lines were treated with control-siRNA or transgene-specific siRNAs. Levels of mRNA corresponding to the transgenes were determined by qPCR and normalised to GAPDH mRNA levels. For each cell lines, the % of mRNA remaining in the transgene-siRNA treated cells as compared to the control-siRNA treated cells are presented (N=2 exp.). A panel of 3 transgenes mutated by targeting the synthetic intron into an exon (light blue label) and 3 transgenes mutated by targeting the synthetic intron into a pre-existing intron (light brown label) were analysed. **(B)** BAC transgenes ESI mutated to be RNAi resistant show RNAi resistance at the protein level. HeLa and clonal HeLa lines expressing ESI-mutated CEP135 or AurkB GFP-tagged BAC transgenes were treated with control, CEP135 or AurkB - siRNAs. Levels of AurkB, CEP135 and Tubulin were monitored by immunoblot using specific antibodies. **(C)** Mutation of GTES1 SxIP motifs into SxNN by ESI mutagenesis disrupts the interaction of a GTSE1-GFP BAC transgene with EB1. Cell lysates from U2OS, U2OS expressing an ESI-mutated GTSE1-GFP SxNN BAC transgene, and U2OS expressing an endogenously GFP-tagged GTSE1 (GTSE1-GFP Cas9), were used in pull-downs with GST or GST-EB1 as baits. Pull-downs inputs and outputs were probed by immunoblot using GTSE1 antibody. GST and GST-EB1 fusion were visualized by Coomassie Blue.

To further demonstrate that ESI-mutagenesis allows the introduction of functional mutations in a BAC transgene, we used the procedure to mutate two EB1-interacting motifs (SxIP) of the microtubule-associated protein GTSE1 to abolish its interaction with the plus end tracking protein EB1(Honnappa *et al.*, 2009; Scolz *et al.*, 2012). In this example, we started with a GFP-tagged GTSE1 BAC, integrated the synthetic intron between the two sites to be mutated, and included the two mutations (SxNN) within the 5’ and 3’ homology arms (Figure S2). U2OS cells were generated stably expressing the BAC transgene and analysed for GTSE1 expression and interaction with EB1. The ESI-mutated BAC yielded a protein of the same size as GTSE1 endogenously tagged with GFP, as expected(Figure 3 C, left panel) (Bendre *et al.*, 2016). However, the mutated protein was no longer efficiently pulled-down by GST-EB1 when compared to the endogenous GTSE1 or GTSE1-GFP (Figure 3 C), verifying that the mutation was present.

### A functional ESI-mutated BAC transgene is expressed in all cells of a clonal line

We showed above that ESI-mutated BACs yield proteins of the expected size and the correct cellular localisation, and that mutation may efficiently confer RNA-resistance or loss of interaction partner when analysed at the cell-population level. We next further assessed whether an ESI-mutated transgene was functionally expressed, i.e. able to sustain the cellular functions of its endogenous counterpart, uniformly in all the cells of a population. This is critical for functional and phenotypic analysis of mutants on the single cell level. To address this question, we ESI-mutated the gene encoding Aurora A kinase (AURKA) on a GFP-tagged BAC transgene to render it resistant to RNAi, transfected it into U2OS cells, and selected a cell clone (U2OS AURKA-GFP siRES (ESI)) that expressed Aurora A-GFP at the same level as the endogenous Aurora A (Figure 4 A, sixth lane). Treatment of this line with siRNA against AURKA led to the depletion of the endogenous Aurora A, while the GFP-tagged Aurora A remained expressed (Figure 4 A, fifth and sixth lanes, lowest and uppermost band respectively) and correctly localised to the centrosomes and the spindle (Figure 4 B and C). U2OS cells depleted of Aurora A accumulate in prometaphase and show smaller bipolar spindles (Marumoto et al JBC 2003, Bird and Hyman 2008) (Figure 4 D, E, and F). In contrast, cells containing the RNAi-resistant Aurora A-GFP ESI maintained a normal distribution of mitotic phases, similar to control-treated cells (Figure 4 D). Furthermore, we could not observe any differences in the length of the metaphase spindle between U2OS treated with control-siRNA and the U2OS AURKA-GFP siRES (ESI) clone treated with the siRNA against AURKA (Figure 4 E and F). This result indicates that all cells depleted for the endogenous Aurora A express a functional ESI mutated Aurora A-GFP ESI transgene. Taken together, these data show that all cells of the U2OS AURKA-GFP siRES (ESI) clone uniformly express a functional Aurora A-GFP protein.

**Figure 4:**
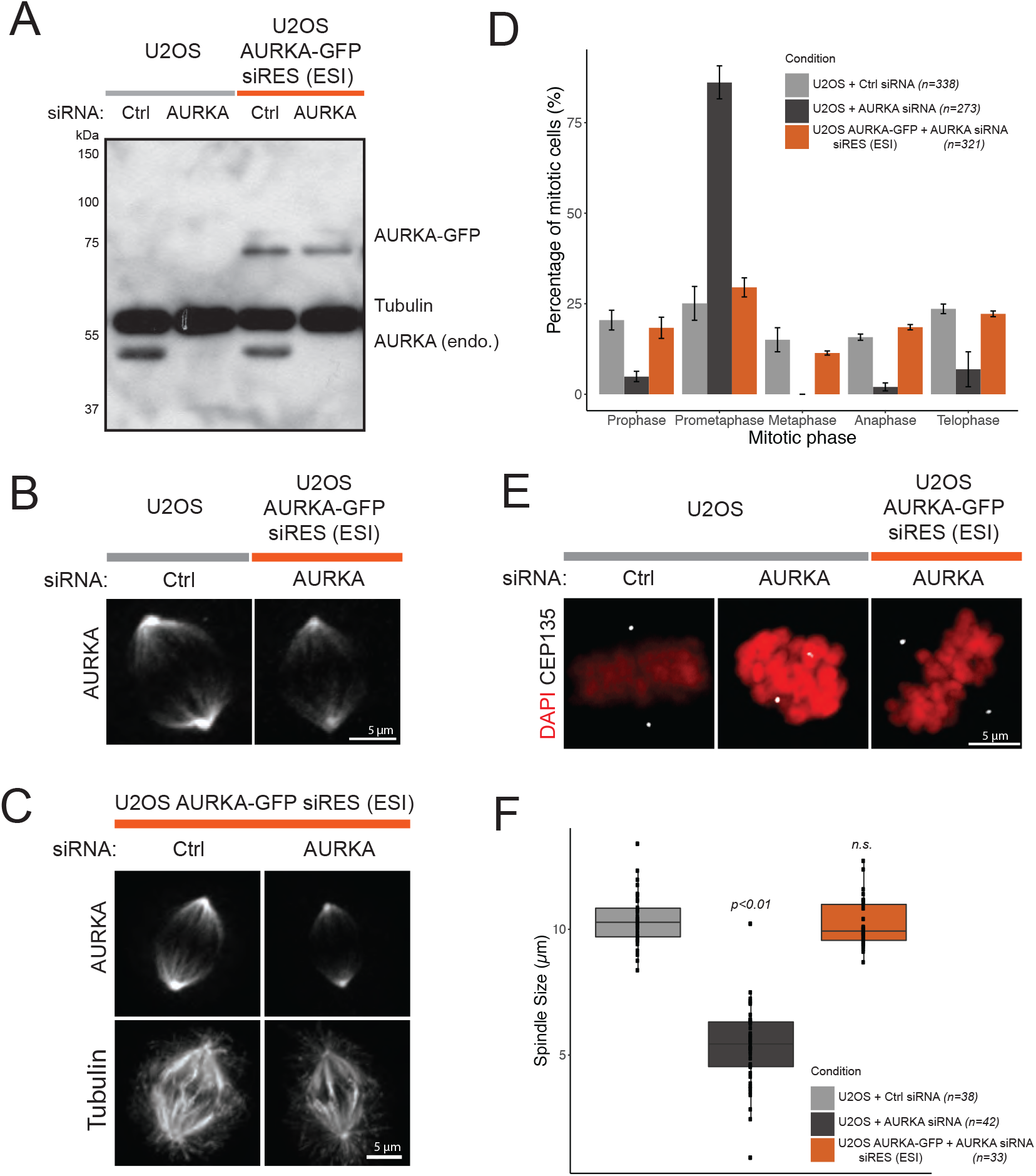
An AURKA-GFP BAC transgene ESI-mutated to carry RNAi resistance rescues endogenous AURKA depletion. **(A)** An AURKA-GFP transgene ESI-mutated to be RNAi resistant is expressed at endogenous level and is RNAi resistant. Immunoblotting on U2OS and a U2OS clone expressing the ESI-mutated AURKA-GFP BAC transgene (U2OS AURKA-GFP siRES (ESI)), both treated with control- or AURKA- siRNA and blocked in mitosis. Antibodies against AURKA and Tubulin were used as probe. **(B,C)** The ESI-mutated AURKA-GFP transgene shows the correct cellular localization. Immunofluorescence on U2OS and U2OS AURKA-GFP siRES (ESI) treated with control- and/or AURKA- siRNA. AURKA antibody (B) or AURKA and Tubulin antibodies (C) were used as a probe. The ESI mutated AURKA-GFP transgene rescues the mitotic arrest **(D)** and spindle collapse **(E,F)** observed after endogenous AURKA depletion. The percentage of cells in each mitotic phase is presented (D) for U2OS and U2OS AURKA-GFP siRES (ESI) clone treated with control- or AURKA- siRNA (N= 3 exp.). U2OS and U2OS AURKA-GFP siRES (ESI) clone were treated with control- or AURKA- siRNA and stained by immunofluorescence using DAPI and an antibody against CEP135 (E). The pole-to-pole distance (μm) in metaphase cells is presented (F). In AURKA-depleted U2OS, spindle length was measured in prometaphase cells with nearly aligned chromosomes. The number of mitotic cells used in each condition (n) and statistically significant differences to U2OS treated with control-siRNA are indicated (Kruskal Wallis, followed by paired Wilcoxon test). All scale bar 5 μm.

## DISCUSSION

We have shown that by co-integrating a selectable marker located within an intron, site-directed mutations can be quickly and efficiently introduced into BAC transgenes using a one-step recombineering procedure. This recombineering method is thus analogous to that for integrating protein tags: a single PCR reaction to generate a cassette with homology arms followed by recombineering and selection. The high efficiency (>90%) of the latter has facilitated high-throughput genome-scale application of protein tagging(Poser *et al.*, 2008; Sarov *et al.*, 2012; Hein *et al.*, 2015; Hasse *et al.*, 2016; Sarov *et al.*, 2016). While we have focused on applications in human cell lines, this method would also be applicable to BAC or fosmids commonly used as transgenes in other model organisms.

Critically, BAC transgenes ESI-mutated to be RNAi resistant yield proteins of the correct size and localization that can functionally rescue the phenotypes observed upon depletion of the endogenous counterpart. Our investigation of transgenic lines at the single-cell level suggest that the synthetic intron is correctly spliced out in all the cells of a population. Furthermore, although we formally describe the ESI-mutagenesis technique here for the first time, we have previously successfully employed it to generate mutations in BAC transgenes, for example, to functionally analyze a microcephaly-associated mutation in the centriolar protein CPAP at the single cell level in HeLa cells and neural progenitor cells(Zheng *et al.*, 2014; Gabriel *et al.*, 2016).

We cannot exclude that a small portion of synthetic introns within individual cells are misspliced, yet without detectable consequences in our functional assays. Indeed, although the core essential sequences for defining introns and splice sites are located primarily within the intron and a few nucleotides into the exons, the precise regulation of efficient splicing is a more complex process and impacted by additional sequence information, including that within exons (Chasin, 2007). Nevertheless, mis-spliced mRNAs are probably minimal and degraded by nonsense-mediated decay.

One of our original concerns in this study was that the new synthetic intron or the gene within may impact expression of the transgene to an unuseful extent. While introns have been well characterized to generally increase gene expression(Nott *et al.*, 2003; Shaul, 2017), nested intronic genes found natively in eukaryotic genomes generally impart a negative effect on the “host” gene expression(Yu *et al.*, 2005; Kumar, 2009). Thus it is possible that expression of a transgene could be impacted by synthetic intron-mutagenesis. While we did not observe gross changes in gene expression in the cell lines generated here, this will likely be highly context-dependent (gene, location of intron, organism, tissue, etc) and we have observed a case where a nested antibiotic resistance gene in a synthetic intron directly integrated into the genome via CAS9 has decreased host gene expression (unpublished results). We have thus also designed a version of the synthetic intron cassette for BAC recombineering in which the antibiotic resistance only carries a bacterial promoter, eliminating the necessity to recombine out with Cre. In the end, because BACs integrate randomly into the genome by standard transfection procedures, clonal variations in expression (+/− 2-fold) in transfected cell lines, likely dependent on position effect and copy number, are anyway observed. Such variations could compensate for synthetic intron effects, and it is thus good practice, prior to phenotypic functional analysis, to select for clonal cell lines expressing BAC transgenes at endogenous levels.

The general strategy outlined here of harbouring a selectable marker gene in an artificial intron adjacent a mutation could in principle be used to increase the efficiency of isolating point mutations via Cas9-induced homologous recombination directly into the genome. This would particularly be useful in hard-to-transfect cell lines where extensive screening may be necessary without a selection step. In our preliminary studies applying this ESI-mutagenesis strategy to direct Cas9-mediated human genome modification, we are indeed able to recover modified cell lines at high frequency. We also found, however, more readily observable impacts on gene expression from the synthetic intron when located in the native gene, which could be alleviated by removing the antibiotic resistance gene after Cre application. While further development of the synthetic intron cassette design/strategy towards direct genome modification holds promise for high efficiency selection of Cas9-mediated site-directed mutation, sensitive applications (i.e. anything to be used in a clinical setting) would not be advisable due to potential changes induced by the synthetic intron.

The use of BAC transgenes remains a powerful genetic tool for exploration of protein function. The ESI mutagenesis method described here allows for straightforward and rapid site-directed mutagenesis of BAC transgenes to further improve their ease of use.

## Methods and Materials

### Bacterial Artificial Chromosomes and recombineering

All BACs used and generated, including modified sequences, are presented in Table S1. LAP- and NLAP-tagged “parental” BACs were generated by recombineering as described in Poser et al 2008. “ESI-mutated” BACs were generated from “parental” BACs following the procedure outlined in Figure S3.

### Antibodies

Mouse anti-GFP (11 814 460 001, Roche), rabbit anti-GTSE1 (custom generated; described in Scolz *et al.*, 2012), and rabbit anti-Aurora kinase B (ab2254, Abcam) were used for Western blots. Mouse anti-α-tubulin (DM1α, Sigma-Aldrich), rabbit anti-Cep135 (custom generated, described in Bird and Hyman, 2008), goat anti-GFP (MPI Dresden; described in Poser *et al.*, 2008), and goat anti-Aurora A (sc-27883, Santa Cruz) were used for both Western Blot and immunofluorescence. Donkey anti-goat Alexa488 (Jackson Immunoresearch; 705 545 147), donkey anti-mouse Alexa594 (Bethyl; A90-337D4), and donkey anti-rabbit Alexa650 (Bethyl; A120-208D5) were used in immunofluorescence. Donkey anti-goat HRP (Santa Cruz; SC-2020), sheep anti-mouse HRP (Amersham; NXA931-1ml), and donkey anti-rabbit HRP (Amersham; NXA934-1ml) were used for Western Blots.

### Cell Lines and Cell Culture

U2OS, HeLa cells, and derivatives were grown at 5% CO2 and 37 °C in DMEM (PAN Biotech) supplemented with 10% filtered Fetal Bovine Serum (FBS, Gibco), 2 mM L-glutamine (PAN Biotech), 0.1 mg/mL streptomycin and 100 U/ml penicillin (PAN Biotech, Pen/Strep mix). BACs were transfected into cells using the Effectene kit (Qiagen) following the manufacturer’s instructions, and stable cell lines were selected on antibiotics.

### RNAi

siRNA sequences used are presented in Table S1. siRNAs were purchased from Thermofisher/Ambion. Unless specified otherwise, control siRNA was Silencer negative control No. 2 (Thermofisher). A reverse transfection approach was used to deplete AurkA and CEP135, using 50 nM and 40 nM siRNA, respectively. The transfection procedure for a 24-well plates with a final volume of 500 μL is described. If required, the procedure was scaled up. The siRNA and 2.5 μL Oligofectamine (Invitrogen) were prepared separately in a final volume of 50 and 15 μL OptiMEM (Invitrogen), respectively. After 5 min incubation at RT, the siRNA and transfection reagent were mixed and further incubated 20 min at RT. Meanwhile, HeLa cells, U2OS cells or their derivatives were seeded at the desired confluency into a 24 well plates containing or not a coverslip. The transfection mix was finally added onto the cells and the volume adjusted to 500 μL with pre-warmed medium. The medium was changed 7 to 8 hours post-transfection. For CEP135 depletion, cells were harvested for Western Blot 48 hours post-transfection. For AurkA depletion, 12 hours post-transfection cells were synchronized with 3.3 μM Nocodazole for 18 hours, and harvested for Western Blot. Alternatively, cells seeded on coverslips were fixed and processed for microscopy 30 hours post-transfection. A forward transfection approach was used to deplete AurkB using 50nM siRNA. Day before transfection, cells were seeded at ~40 % confluency in a 3.5 cm dish. The siRNA and 5 μL of Oligofectamine (Invitrogen) were prepared separately into 200 μL OPTIMEM. After 5 min incubation at RT siRNA and Oligofectamine solutions were mixed and further incubated 20 min at RT. Meanwhile, cells were washed twice with PBS and 1.6 mL OPTIMEM were added on top of them. The transfection mix was then added onto the cells. Cells were harvested for Western Blot 48 hours post-transfection. In experiments measuring mRNA levels in HeLa cells, a control scrambled RNA (customed, Ambion, sense: 5’-UUCUCCGAACGUGUCACGUtt-3’) was used, and siRNA was transfected using Interferin (Polyplus Transfection) and 100nM siRNA per well of a 6-well plate in forward or reverse transfection (140,000 cells). Duration of the depletion is indicated for each gene in Table S1.

### Western Blot

Cells were lysed on ice for 10 to 15 min in RIPA buffer or cell lysis buffer (50 mM Na2HPO4, 150 mM NaCl, 10% glycerol, 1% Triton X-100, 1 mM EGTA, 1.5 mM MgCl2, 50 mM HEPES pH 7.2, 1 mM DTE) both supplemented with 2X protease inhibitor mix (SERVA). Debris were eliminated by centrifugation at 4 °C at max-speed in a benchtop centrifuge. Samples were prepared in Laemmli buffer and run on SDS-PAGE before transfer onto a nitrocellulose membrane. 5% milk (Carl Roth) in PBS Tween20 0.1% (SERVA electrophoresis) was used to block the membrane before incubation with the primary and secondary antibodies diluted in the same blocking solution. Membranes were incubated with ECL reagent (GE Healthcare) before development onto Hyperfilm (Amersham) or imaging on the ChemiDoc™ MP imaging system (BioRad). Digital data were obtained by scanning Hyperfilms or using the ImageLab software (BioRad). Fiji was used to adjust levels, generate 8-bit tiffs, and crop images.

### qPCR analysis

cDNA was isolated from 250,000 cells per condition using the Cells-to-cDNA II Kit (Ambion AM1722) following the manufacturer’s two-step RT-PCR protocol. qPCR was performed with SYBR Green (ThermoFisher) on a Mx3000 qPCR Machine (Stratagene). Primers are indicated in Table S1.

### Pull-downs

Cells from a confluent 15 cm dish were lysed for 15 min on ice in 2mL cell lysis buffer (50 mM Na2HPO4, 150 mM NaCl, 10% glycerol, 1% Triton X-100, 1 mM EGTA, 1.5 mM MgCl2, 50 mM HEPES pH 7.2, 1 mM DTE) supplemented with 2X protease inhibitor mix (SERVA) and 1mM DTE (15min on ice). Debris were removed by centrifugation at max-speed at 4 °C in a benchtop centrifuge. Total protein concentration was measured by Bradford. Equivalent amounts of cell lysate were incubated 20 hours at 4 °C on GST or GST-EB1 beads. Beads were washed thrice with 1 mL cell lysis buffer supplemented with 2X Protease inhibitor. Proteins were eluted with 45 μL Laemmli 2x.

### Immunofluorescence

For most experiments, cells were seeded onto coverslips and fixed in ice-cold methanol for 12 min at −20 °C. Cells were blocked at RT with 5% BSA (Sigma) dissolved into PBS. For images of Aurora B-GFP and CHMP4B-GFP cell lines, cells were fixed in 3% PFA supplemented with 5 mM EGTA, 1mM MgCl2 and 2% sucrose, they were washed, then permeabilized using PBS/0.1% Triton X-100 and eventually blocked with PBS/0.2% Fish Skin Gelatin (Sigma, Germany). Primary antibodies were diluted into 5% BSA in PBS and incubated onto coverslips for 1 h at 37 °C in a wet chamber. The same procedure was applied for secondary antibodies. Coverslips were mounted using Prolong Gold antifade mounting with DAPI (Molecular Probes and Thermo Fisher Scientific). Cells were washed thrice with PBS between every step.

### Microscopy

Immunofluoresence images for the AurkA experiments were acquired using a spinning-disk confocal system (Marianas, Intelligent Imaging Innovation, 3i) built on an Axio Observer Z1 microscope (Zeiss) with an ORCA-Flash 4.0 camera (Hamamatsu Photonics). Images were taken with a 63× 1.4 NA Plan-Apochromat oil objective (ZEISS). The spindle size was measured using the CEP135 signal in the Slidebook software (Marianas, Intelligent Imaging Innovation, 3i, V5.5 or later).

Experiments assessing the localization of the GFP transgenes, were all performed on live cells at the exception of the AurkB and CHMP4B transgenes. Live cells were imaged in CO2-independent visualization media (Gibco). Images were acquired using a DeltaVision imaging system (GE Healthcare) with an sCMOS camera (PCO Edge 5.5) and a 60x 1.42NA Plan Apo N UIS2 objective (Olympus).

---

**Figure S1:**
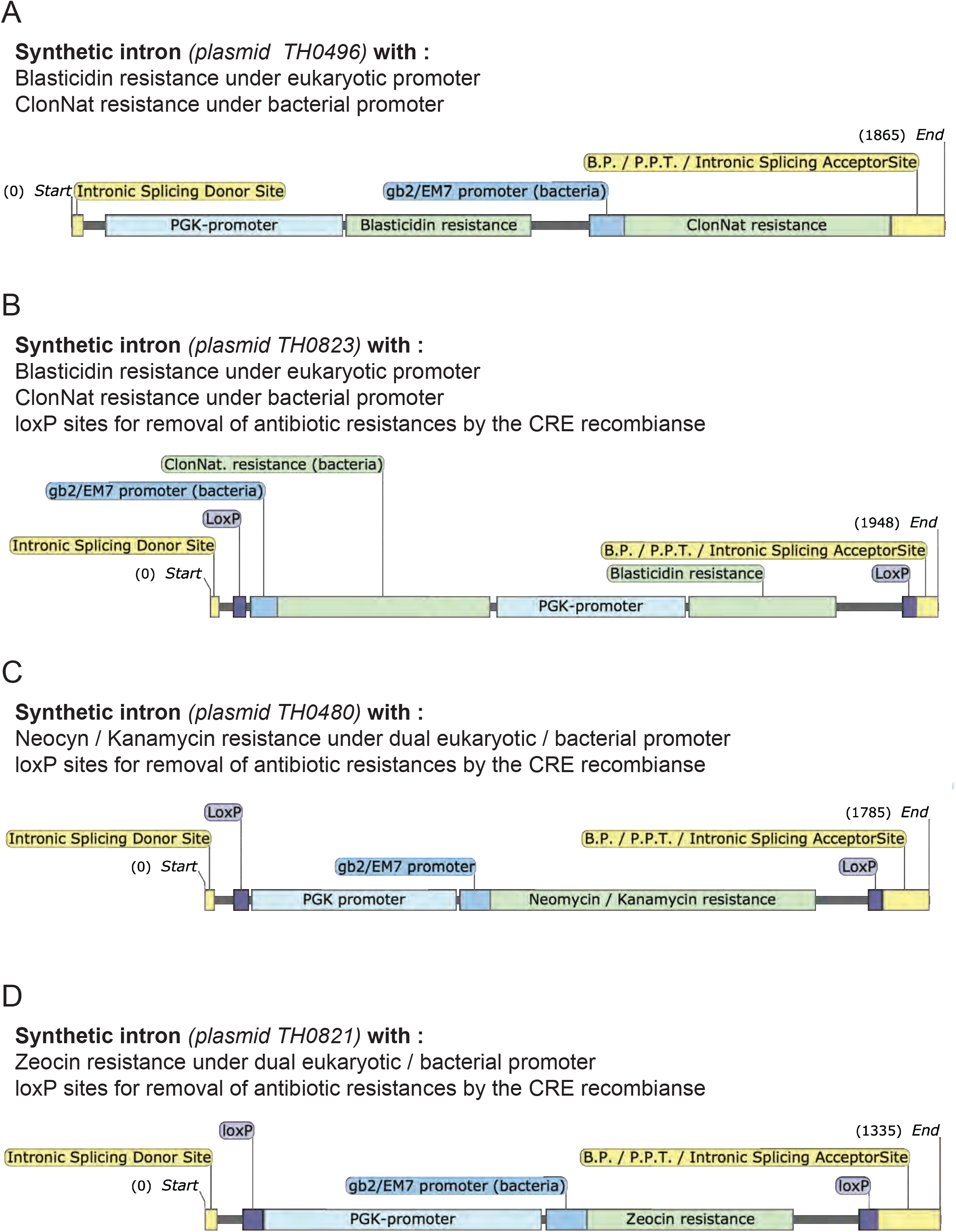
Available synthetic intron cassettes. The synthetic intron cassettes generated in this study are depicted (A to D). The resistance genes and the promoters (bacterial and eukaryotic) controlling their expression are shown. When present, the loxP sites allow the removal of the antibiotic resistances by the CRE recombinase. The position of the core intron-based splicing signal sequences (intronic splicing donor dite, branch point (B.P.), polypirimidine tract (PPT), intronic splicing acceptor site) is shown. The plasmid from which each cassette can be amplified is indicated. Numbers indicate the nucleotide positions within the cassette.

**Figure S2:**
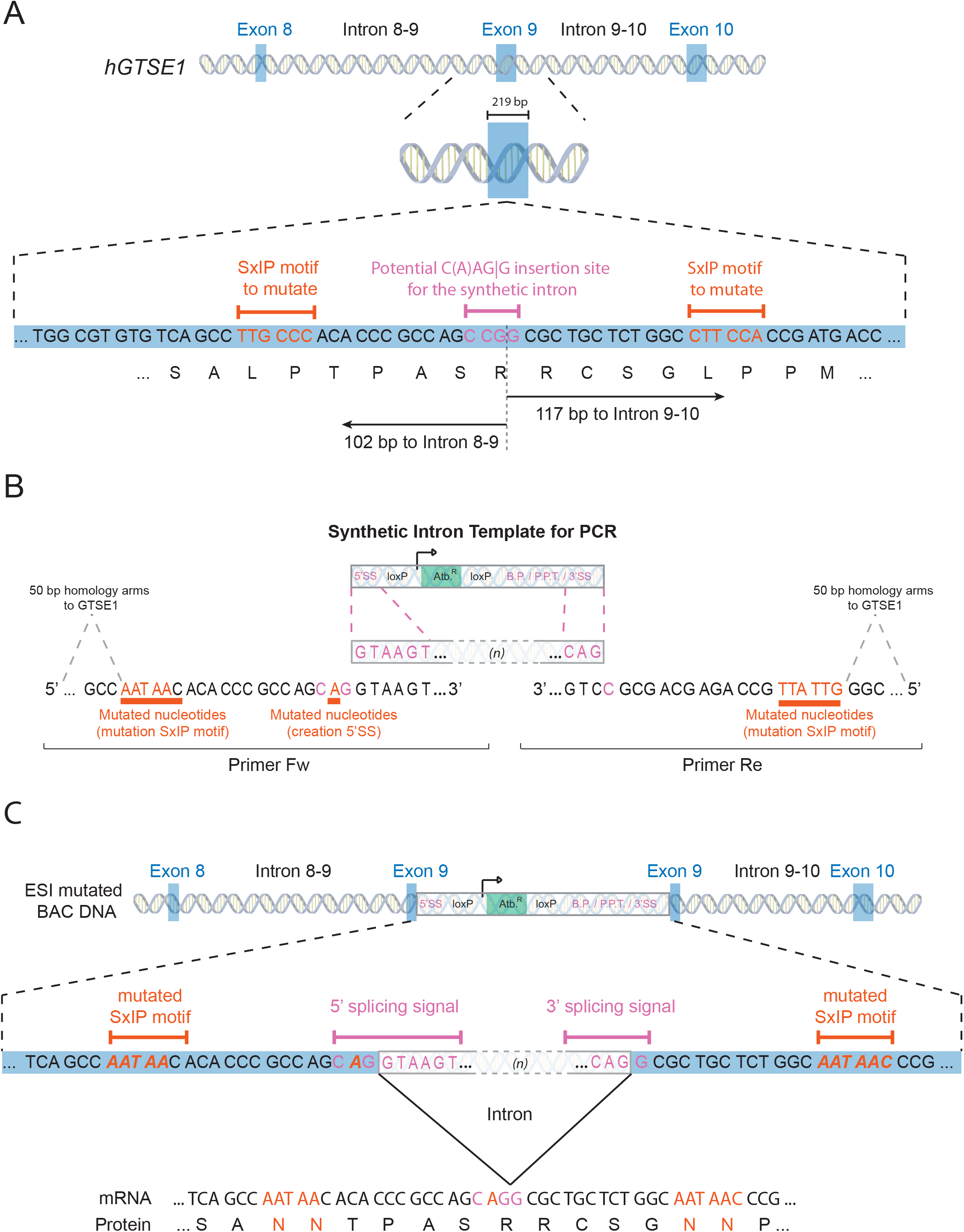
ESI mutagenesis of the GTSE1 SxIP motifs. **(A)** Scheme representing the sequence of exon 9 of GTSE1 containing the two SxIP motifs (in orange) and a potential insertion site for the synthetic intron (in pink). Distance (in bp) of the potential insertion site for the synthetic intron to the border with introns 8-9 and 9-10 are presented. **(B)** Primers used to amplify the synthetic intron are presented. They are composed of (i) homology arms to GTSE1, (ii) the SxIP mutations (in orange) and (iii) a synonymous mutation (in orange) allowing the potential insertion site (in pink) to fit the CAGG consensus. **(C)** The amplified synthetic intron is recombined onto the GTSE1 BAC. The synthetic intron is spliced out from the pre-mRNA to yield a mRNA coding GTSE1 mutated at both SxIP motifs.

**Figure S3:**
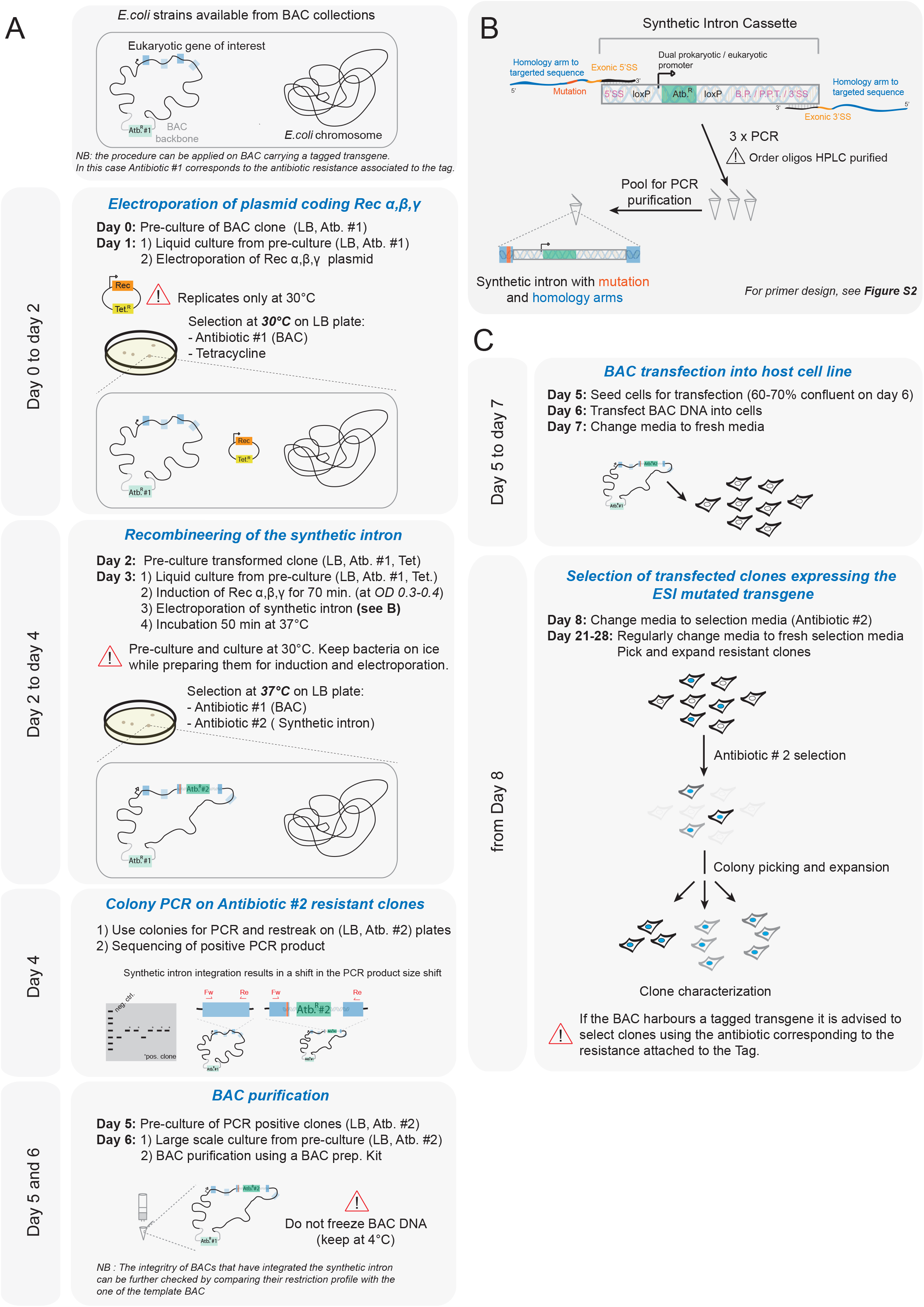
The ESI mutagenesis procedure at a glance. (A) Scheme presenting the BAC recombineering protocol. Important parameters to pay attention to are indicated. (B) Scheme showing the procedure to obtain the ESI mutagenesis BAC recombineering template. Three PCRs (see Figure 1B and S1 for primer design) are performed to amplify the synthetic intron and pooled for subsequent PCR purification. The resulting product is used as recombineering template. (C) The ESI mutated BAC is transfected into eukaryotic cells. Stably transfected cells are selected using the antibiotic resistance coded by the synthetic intron. If the BAC carries a fluorescent protein tagged transgene, the antibiotic resistance associated to the tag should be use for selection.

## Notes

### Competing Interest Statement

The authors have declared no competing interest.

